# Word structure tunes electrophysiological and hemodynamic responses in the frontal cortex

**DOI:** 10.1101/2021.11.14.468513

**Authors:** Fei Gao, Lin Hua, Yuwen He, Zhen Yuan

## Abstract

To date, it is still unclear how word structure might impact lexical processing in the brain for morphological impoverished language like Chinese. In this study, concurrent EEG and fNIRS recordings were performed to inspect the temporal and spatial brain activity that are related to the morphological priming effect (compound/derivation constitute priming vs. non-morphological priming) and word structure (compound vs. derivation) modulation. Interestingly, it was discovered that the morphological priming effect was mainly detected by the behavioral performance and spatial brain activation in the left prefrontal cortex, while word structure effect was revealed by the behavioral data as well as the temporal and spatial brain activation patterns. In addition, Chinese derivations exhibited significantly enhanced brain activation in the frontal cortex and involved more brain networks as compared to lexicalized compounds. The results were interpreted by the differing connection patterns between constitute morphemes within a given word structure from spreading activation perspective. More importantly, we demonstrated that Chinese word structure effect showed a distinct brain activation pattern as compared to that from the dual-route mechanism in alphabetic languages. Therefore, this pilot work paves a new avenue for comprehensively understanding the underlying cognitive neural mechanism associated with Chinese derivations and coordinate compounds.

## Introduction

In linguistics, morphology concerns not only how words are formed, but also how they are inter-connected with each other in the arguable mental lexicon (Levelt, 1993; Treisman, 1960). With developments in experimental psychology and neurosciences, psycholinguistic and neurolinguistic approaches in studying morphology in the human mind/brain have provided accumulating evidence for sub-lexical information processing and morphological structure processing. For instance, a key issue pertaining to morphological processing in the past five decades discussed about whether words are stored and processed in a full-listing or full-parsing manner for morphologically complex words (De Grauwe, Lemhöfer, Willems, & Schriefers, 2014; Frost, Kugler, Deutsch, & Forster, 2005; Fei Gao, Wang, Zhao, & Yuan, 2021), which yet has not reached consensus (Jiang, 2018; Schiller & Lieber, 2020). Importantly, irrespective of whether decomposition is an obligatory process of word recognition, the way how morphemes/lexemes are connected to establish a new word has been recognized to significantly modulate lexical access. Therefore, the aim of this study is to looks into the neural underpinnings of morphological structure processing of Chinese words.

To shed light on the sub-types of word morphological structures, neuroimaging and behavioral studies have been carried out to inspect the representations of inflectional and derivational words in alphabetical languages (Bölte, Jansma, Zilverstand, & Zwitserlood, 2009; Carrasco-Ortiz & Frenck-Mestre, 2014; Grainger & Beyersmann, 2020; Leminen, Smolka, Dunabeitia, & Pliatsikas, 2019; Whiting, Marslen-Wilson, & Shtyrov, 2013). Inflections are used upon word forms to mark grammatical changes like gender, number, voice, and tense, whose meanings remain unchanged. By contrast, derivations are characterized by attaching affixes (prefixes, infixes, and suffixes) to base words, thus establishing new word form, meaning, and even grammatical category. For example, “*walks*” in “*Tom walks to school every day*” is an inflection marking third-person singular verb in present tense, while “*walker*” is a noun derived from the verb “*walk*”. Specifically, Newman, Ullman, Pancheva, Waligura, and Neville (2007) compared the event-related potential (ERP) responses between the regular/irregular past tense of English verbs, phrase structure, and lexical semantics in sentential contexts. It was discovered that irregular past tense elicited significant left-lateralized anterior negativities (LANs), probably indexing a rule-based computation on morphology, while significant P600 effects identified from regular and irregular violations might suggest a controlled processing for lexicalized linguistic items (e.g., irregular forms like *ran*). This pattern was replicated in a recent Swedish study (Schremm, Novén, Horne, & Roll, 2019) and elaborated in a so-called dual-access model (Baayen, Dijkstra, & Schreuder, 1997; Bozic & Marslen-Wilson, 2010; Marslen-Wilson & Tyler, 2007) highlighting the neurocognitive signatures for morphological complexity. In this dual system, the processing of regular inflections in English might selectively engage the left fronto-temporal network, which is sensitive to rule-based decomposition and combination. By contrast, the bilateral subsystem and broader brain regions are employed in reading derivational words and highly lexicalized forms, which is neuro-biologically specified for whole-word and storage-based sound-to-meaning mapping.

In addition, compared to inflections and derivations, few studies have been performed to examine the compound processing and the sub-structures of compounding. Compounding involves the process of concatenating different lexical constitutes (words/lexemes) to create new lexical items, which might represent a fundamental mechanism of morphological productivity for most languages (Libben, 2014; Libben, Gagné, & Dressler, 2020). The majority of existing studies focused on how constitute frequency (e.g., Andrews, Miller, & Rayner, 2004), semantic transparency (e.g., Brooks & Cid de Garcia, 2015; Libben, Gibson, Yoon, & Sandra, 2003; Smolka & Libben, 2017) and constitute headedness (e.g., Libben et al., 2003; Marelli & Luzzatti, 2012) would impact lexical access and morphological parsing in compound word recognition (Leminen et al., 2019). Now what remains unclear is the extent to which word structure (i.e., the grammatical and semantic relations between lexical constitutes) would modulate the representation and processing of compound words. Insights from conventional descriptive linguistics (see review by Olsen, 2012) suggested a variety of compound structures, partially based on syntactic relations yet mostly on semantics, including apposition (e.g., *woman doctor*), subject/object + action (e.g., *sunrise*), and purpose (e.g., *wineglass*), and others (Jespersen, 1914). However, present psycholinguistic studies on compound processing exclusively focused on “modifier-head” (e.g., teacup, pineapple) structure (Brooks & Cid de Garcia, 2015; Smolka & Libben, 2017). Although a couple of studies mentioned the existence of coordinative structure in English, or so-called “dual compounds”/ “copulative type” (e.g., *singer-songwriter, architect-sculptor, in-and-out*) that is primarily a hyphenated combination (Chung, Tong, Liu, McBride-Chang, & Meng, 2010; Libben, 2014; Olsen, 2012), no study has been conducted to inspect the neural basis of this structure or make the comparisons between various compounding structures. The inadequate evidence on this issue might be attributable to the relative impoverished compounding system of English words.

More interestingly, the Chinese language is categorized as one without much inflection (see the debate about Chinese inflection from Dai, 1997), which is in striking contrast to alphabetical languages. According to structuralism linguistics (e.g., B. Huang & Liao, 2011), Chinese vocabulary is classified into four categories in terms of word structure including mono-morphemic word (e.g., *葡萄*, pu2-tao, “grape”), derivation (prefixed and suffixed), reduplication (e.g., *妈妈*, ma1-ma1, mom-mom, “mother”), and compound, among which compound words constitute the largest portion (more than 70%). According to a survey through modern Chinese dictionary (Cao, 2003), the majority of multi-morphemic words in colloquial dataset is compound words (83.02%), followed by derivations (14.51%). In particular, there are five sub-structures of compounds, including modification (54%, e.g., *黑板*, hei1-ban3, black-board, “black-board”), coordination (26%, e.g., *花草*, hua1-cao3, flower-grass, “plant”), verb-object (18%, e.g., *吃饭*, chi1-fan4, eat-food, “to eat”), verb-resultative (2%, e.g., *长大*, zhang3-da4, grow-up, “grow up”), and subject-predicate (1%, e.g., *晚安*, wan3-an1, night-safe, “good night”) (statistics from Su, 2016). The seminal work concerning Chinese word structure effect on morphological decomposition revealed that compounding structure (coordination vs. modifier) might modulate the constitute frequency effect in word recognition (B. Zhang & Peng, 1992). Specifically, it was discovered that the frequency of both constitutes in coordinative structure impacted the reaction time (RT) to target words, whereas for modifier structure, RT was only related to the constitute on the final position. These findings shed light on the decomposition mechanism of Chinese compound words, which is also mediated by morphological structures. More importantly, both behavioral and neuro-imaging studies since then have accumulated ample evidence for the cognitive neural mechanism associated with the morphemic effect of Chinese word reading. For example, ambiguity and polysemant at the constitute morpheme level would significantly affect the representation of disyllabic compound words (e.g., C. Y. Huang, Lee, Huang, & Chou, 2011; H. W. Huang, Lee, Tsai, & Tzeng, 2011; L. Liu et al., 2013; Tsang, Wong, Huang, & Chen, 2013; Wu, Duan, Zhao, & Tsang, 2020; Zhao, Wu, Tsang, Sui, & Zhu, 2021; Zou, Packard, Xia, Liu, & Shu, 2015, 2019). In addition to the morphemic effect, morpheme relations were also investigated in terms of semantic/thematic (Ji & Gagné, 2007; Jia, Wang, Zhang, & Zhang, 2013; Li & Xu, 2021; Pan & Jared, 2020; Z. Xu & Liu, 2019) and grammatical aspects.

In addition, existing behavioral and electrophysiological studies mostly concentrated on subordinate and coordinate compounding structures and their corresponding representational difference in mental lexicon. Drawing on a compounding production task, P. D. Liu and McBride-Chang (2010b) compared the Chinese third graders’ word production performance with four compounding structures. Their results reveled that subordinate and coordinate structures were easier to produce than subject–predicate and verb–object structures, whose difficulty was detected to be proportional to their distributional characteristics in language units (Laudanna & Burani, 1995). Further studies differentiated the subordinate and coordinate structures with various paradigms. For example, subordinate structure was harder to be recognized than coordination in a priming lexical decision task, where semantic relatedness and structural consistency between primes and targets were manipulated (P. D. Liu & McBride-Chang, 2010a). In subordinate condition, the same structures facilitated the semantic priming effect while the coordinate structures manifested the opposite pattern. Meanwhile, subordinate structures were detected to significantly boost literacy performance in memorizing compound words (D. Liu, 2016, 2017), relative to coordinative structures. D. Liu (2016) attributed this difference to the strengths of constitute connections within a compound word in light of the spreading activation theory (Collins & Loftus, 1975) and further proposed the independent representation of morphological structure in mental lexicon (D. Liu, 2017). Both of the two constitute morphemes in a coordinate structure contribute equally to the whole word meaning (e.g., *风雨*, feng1yu3, wind-rain, “storm”), whose inter-connection are relatively weaker than the modifier-head relation in subordinate structure. It therefore requires more cognitive efforts to combine the lexical constitutes for coordinate structures. Importantly, there might exist a morphological structure layer in mental lexicon (D. Liu, 2017), in addition to morpheme and word layers in mental lexicon, which is originally proposed by the interactive activation model (Taft, 1994). At this specific layer for word structures, both the semantic and syntactic connections between lexical constitutes would be activated. Given the relative weak semantic connections in coordinate structure relative to subordinate structure, its corresponding morphological effect might be dampened.

However, the different patterns between coordinate and subordinate structures from Liu et al. were not replicated in Cui et al. (2018) and Chung et al. (2010). The inconsistency was thought to result from the prolonged stimulus onset asynchrony (SOA=200 ms). For Chung et al.’s ERP study, only coordinate compounds was used, and morphological structure effect was null in behavioral data although manifested by the P250 effect (220-300 ms time windows). Structural priming elicited greater P250 than distinct structure pairs, which might indicate a word structure facilitation and top-down processing at the early stage of lexical access (Hill, Strube, Roesch-Ely, & Weisbrod, 2002). The discrepancy calls for further examinations concerning the morphological structure effect in coordinate compounds.

Only a handful of neuro-imaging studies quantified the brain changes associated with Chinese morphological processing (F. Gao et al., under review; Ip, Hsu, Arredondo, Tardif, & Kovelman, 2017; Ip et al., 2019; Zou et al., 2015). They identified that both the left frontal and temporal cortex are involved in processing Chinese morphology, yet failed to differentiate various structures (i.e., mixed with different sub-structures of compounding and derivations). To our knowledge, there is only one functional neuroimaging study which tried to disassociate the brain responses of various sub-structures of Chinese compound words (Hsu, Pylkkänen, & Lee, 2019). Hsu and colleges examined the Chinese morphological complexity effect (mono-morphemic vs. multi-morphemic) and compounding structural effect (subordinate, coordinate, and verb-object) by using lexical decision task and magnetoencephalography (MEG) technique. It was discovered that compound words generated greater brain activations at an early stage (200 ms) in the left temporal cortex. In the later time window of 300-400 ms, however, there was a null effect in coordinate structure as compared to that from the baseline (mono-morphemic word), while both subordinate and verb-object structures generated greater responses in the posterior part of the left temporal region. This work attributed the absence of coordinate structure effect in the left temporal cortex to the fact that coordinate compound words might lack a specifier-head-complement relational structures as manifested in subordinate and verb-object words. Instead, the two component morphemes make equal contributions to the whole-word meaning, resulting in a rather loose connection. However, this study failed to include the frontal cortex as a region of interest (ROI) in brain activation analysis, while related studies revealed that the left frontal cortex is also a crucial hub for Chinese morphological processing (F. Gao et al., under review; Ip et al., 2017), indicating it is essential to examine the role of frontal cortex in processing different compound sub-types.

Therefore, the neurobiological basis of differing Chinese compound structures is still poorly understood, with respect to both temporal signatures and localizations in the human brain. Specially, much uncertainty still exists about the neural reality of coordinate structure effect. Importantly, no previous study has investigated the brain representations of Chinese derivation words, even though they constitute the second largest proportion of Chinese vocabulary. The present study therefore aims at exploring whether Chinese derivations use the similar cognitive resources as compound words and examining the root priming effects from derivations and compounds in relation to semantic controls (non-morphological relationship). To address this issue, we will inspect what the temporal and spatial specifications in the human brain are associated with the morphological priming effect (compound/derivation constitute priming vs. non-morphological priming) in Chinese word reading. In addition, we will also inspect whether Chinese derivational word reading utilizes the same neural resources as Chinese compound words (derivation vs. compound). Consequently, masked priming is used to elicit early and automatic morphological parsing (Fiorentino, Naito-Billen, Bost, & Fund-Reznicek, 2014) in a visual lexical decision task. Electroencephalogram (EEG) and functional near-infrared spectroscopy (fNIRS) were recorded simultaneously to depict both temporal signatures and spatial activations in the brain, as the latter one would somewhat compensate for the relatively poor spatial resolution of EEG.

## Materials and Methods

### Participants

Thirty Mandarin Chinese native speakers (mean age: 22.2 ± 3.2 years old; 15 female) were recruited from the University of Macau campus. They were registered university students from various majors. All participants were right-handed and reported no neurological illness or mental disorder, with normal or corrected-to-normal vision. All materials and procedures were approved by the Institutional Review Board at the University of Macau. Written consent form was obtained from each participant prior to the experiment.

### Stimuli materials

Thirty-six disyllabic Chinese compound words with coordinate structures and 36 disyllabic derivational words with suffixes (e.g., *儿*/er/, *民*/min2/) were selected as target words. In masked priming task (Lavric, Clapp, & Rastle, 2007), these words were primed by their corresponding constitute morpheme on the head position. In addition, another 36 disyllabic compound words with four lexical structures (subordinate: 22, coordinate: 8, verb–object: 5, noun-complement: 1) were selected as control group with a semantic related Chinese character as the prime. In control group, prime character and constitutes of target word are distinct at both orthographic and phonological levels and yielded no morphological relationship. Twelve Chinese native speakers were invited to rate the semantic relatedness between the prime and target among 36 derivation, 36 compound and 36 control materials with a 0-7 Likert scale. The averaged semantic relatedness from each group was above five, indicating a high semantic transparency and consistency among groups. In particular, the three groups of words (three lexical conditions) are matched regarding the word frequency, character frequency, and number of strokes (Table 1).

**Table 1.** Stimulus statistics across three lexical conditions.

|  | Prime | Frequency | Stroke Number | Target | Frequency | Stroke Number | Cloze Probability | Semantic Relatedness |
| --- | --- | --- | --- | --- | --- | --- | --- | --- |
| Derivational word | 网/wang3/, net | 94 (145) | 8.3 (2.3) | 网民/wang3 min2/, netizen | 8 (32) | 15.3 (3.8) | 0.07 (0.12) | 5.2 (0.9) |
| Compound word | 花/hua1/, flower | 83 (109) | 8.6 (3.1) | 花草/hua2 cao3/, plant | 7 (10) | 16.9 (4.6) | 0.12 (0.19) | 5.4 (0.9) |
| Non-morphological | 脏/zang1/, dirty | 356 (1076) | 8.8 (2.8) | 污水/wu1 shui3/, dirty water | 34 (79) | 17.8 (4.5) | / | 5.3 (0.5) |

Besides, 108 disyllabic Chinese non-words were selected from MELD-SCH dataset (Tsang et al., 2018) as the fouth non-words condition. The non-words yielded a pronounceable meaningless string and are primed by a distinct character. They worked as fillers to balance the yes/no responses.

### Procedures

The visual lexical decision task was presented for this study, which was adopted from the morphological priming paradigm with a short stimulus-onset asynchrony (SOA) (Chung et al., 2010) and a masked priming technique (Lavric et al., 2007). E-prime was used to program the materials and procedures (Figure 1), in which a white fixation was first presented in the screen center of a PC for 300 ms, followed by a blank of 200 ms. And then a series of asterisks serving as the mask were presented for 300 ms and subsequently would be replaced by the prime in 57 ms. After the prime disappears, participants would read the target and decide whether the displayed character string is a real Chinese word or not by pressing the corresponding buttons labelled as “yes” or “no” in the keyboard as quickly as possible. If they fail to react within three seconds, it would be marked as wrong response automatically by the task. After the response, there would be blank jittered from one to four seconds (P. Zhang, Zhang, Peng, Song, & Bai, 2016). The real Chinese word case consists of the three lexical conditions. Noted here that primes were presented with Italic *Kaiti* typeface and size of 40, whereas targets were displayed with bold *SimHei* at 40. Ten practice trails with correctness feedback would be offered before the formal experimental tests.

**Figure 1.**
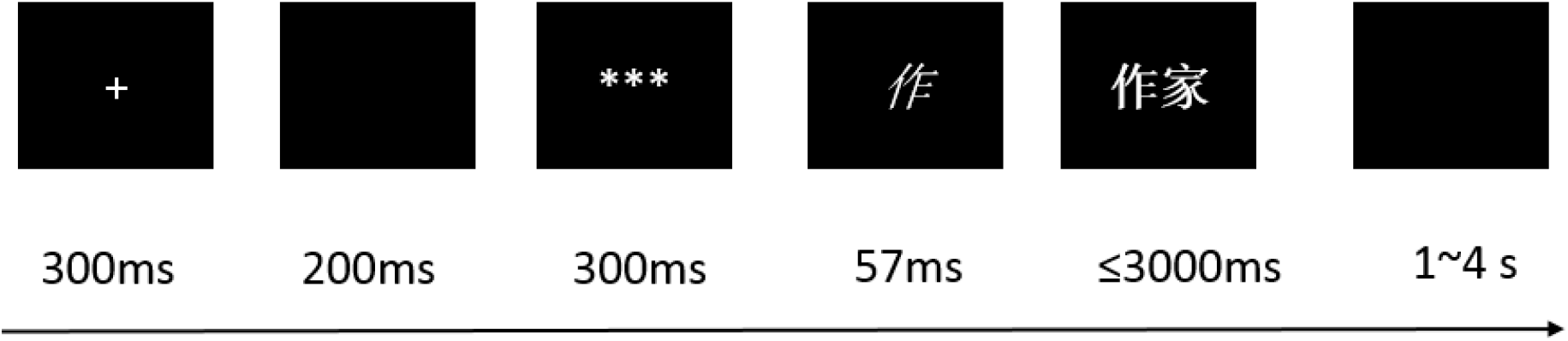
Schematic of the lexical decision task. Participants need to decide whether the target is a real word or not when they read the character string. The stimulus-onset asynchrony (SOA) between prime and target is set at 57 ms in light of the previous study (Chung et al., 2010). “*作*” denotes “write” and “作家” stands for “writer”.

### EEG recordings and data analysis

EEG and fNIRS data were collected simultaneously using an EasyCap (Brain Products, Munich, Germany), which was connected to Brain Products EEG system and NIRScout system (NIRx Medizintechinik GmbH, Berlin, Germany). For EEG acquisition, 32 electrodes were arranged on the cap based on the international 10/20 system. EEG data were digitized at 500 Hz with a bandpass filter of 0.03-70 Hz. During the online acquisition, the left mastoid electrode was used as a reference and impedance of all electrodes was kept below 20 kΩ.

For offline analysis in EEGLAB v2021.0, continuous EEG data were firstly re-referenced to the grand average of all channels and filtered with a band pass of 1-30 Hz. Bad channels were interpolated by averaging the spherical electrodes, which took up less than 3% of all channels. And then data segmentation was completed, which consisted of 158 ms before the target onset and 1000 ms afterwards. Eye movement components were removed from the segmented data by Independent Component Analysis (ICA) algorithm and ICLabel plugin (Pion-Tonachini, Kreutz-Delgado, & Makeig, 2019). Bad epochs were further removed by visual inspection.

The averaged amplitudes of selected electrodes were computed and compared in the time windows of 220-300 ms (P250) and 300-500 ms (N400) according to the detected ERP components (Chung et al., 2010). Specifically, AFF5h, FC1, FCz, FC2, and FC6 from the bilateral and midline sites of frontal cortex were examined for the ERP component P250, while centro-parietal electrodes CPP5h, CP1, CP2, and CP6 were inspected for the N400 effect.

### fNIRS recording and data analysis

Eight LED light sources and 8 detectors were placed in the left frontal and temporal cortex (P. Zhang et al., 2016), thus generating 22 fNIRS channels (Figures 2B and 5A). Each source transmitted LED lights at the wavelength of 760 nm and 850 nm. The distance between each light source and detector was 3 cm. Optical signals were acquired at the sampling rate of 7.81 Hz. The MNI coordinates of all optodes and channels were obtained from their spatial information at the international 10/20 system, which were then imported into NIRS_SPM software (Ye, Tak, Jang, Jung, & Jang, 2009) to generate anatomical label and percentage of overlap.

**Figure 2.**
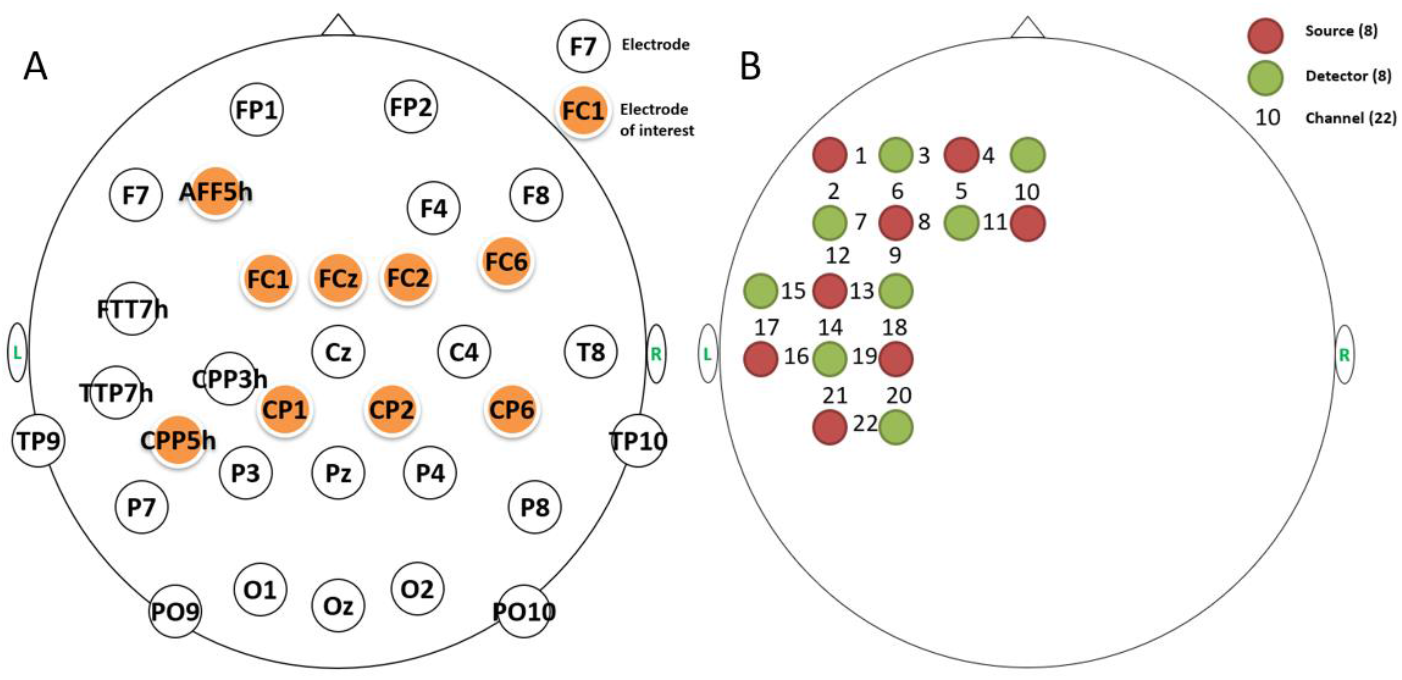
The layout of fused EEG-fNIRS arragement. (A) There were 32 electrodes in total, nine of which (in orange) were used for further ERP analysis. (B) Eight light sources and 8 detectors generated 22 fNIRS channels, covering the frontal and temporal regions of the left hemisphere.

**Figure 3.**
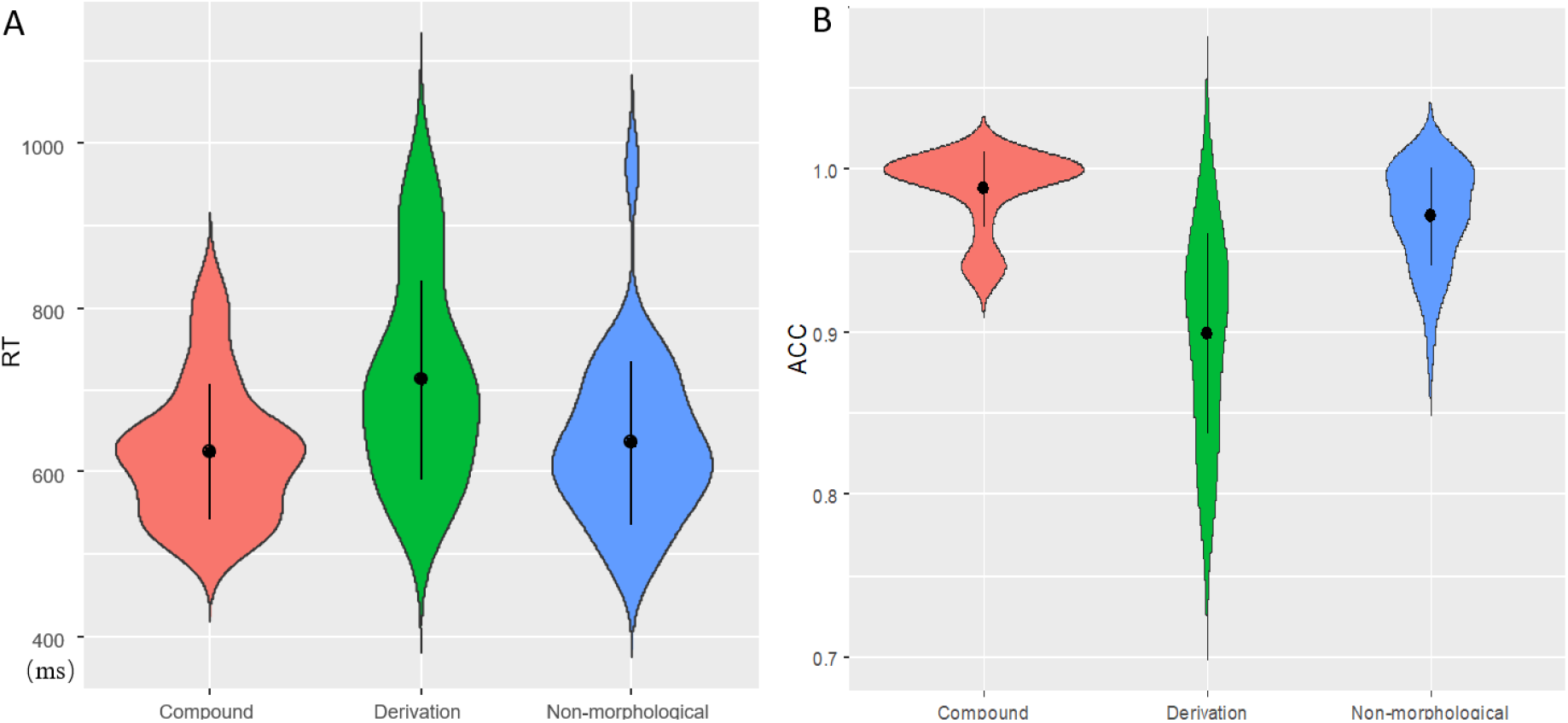
Violin plots of behavioral results. Black points denote means, whereas the error bars define standard deviations. RT differences (A), and ACC difference (B) across three conditions.

**Figure 4.**
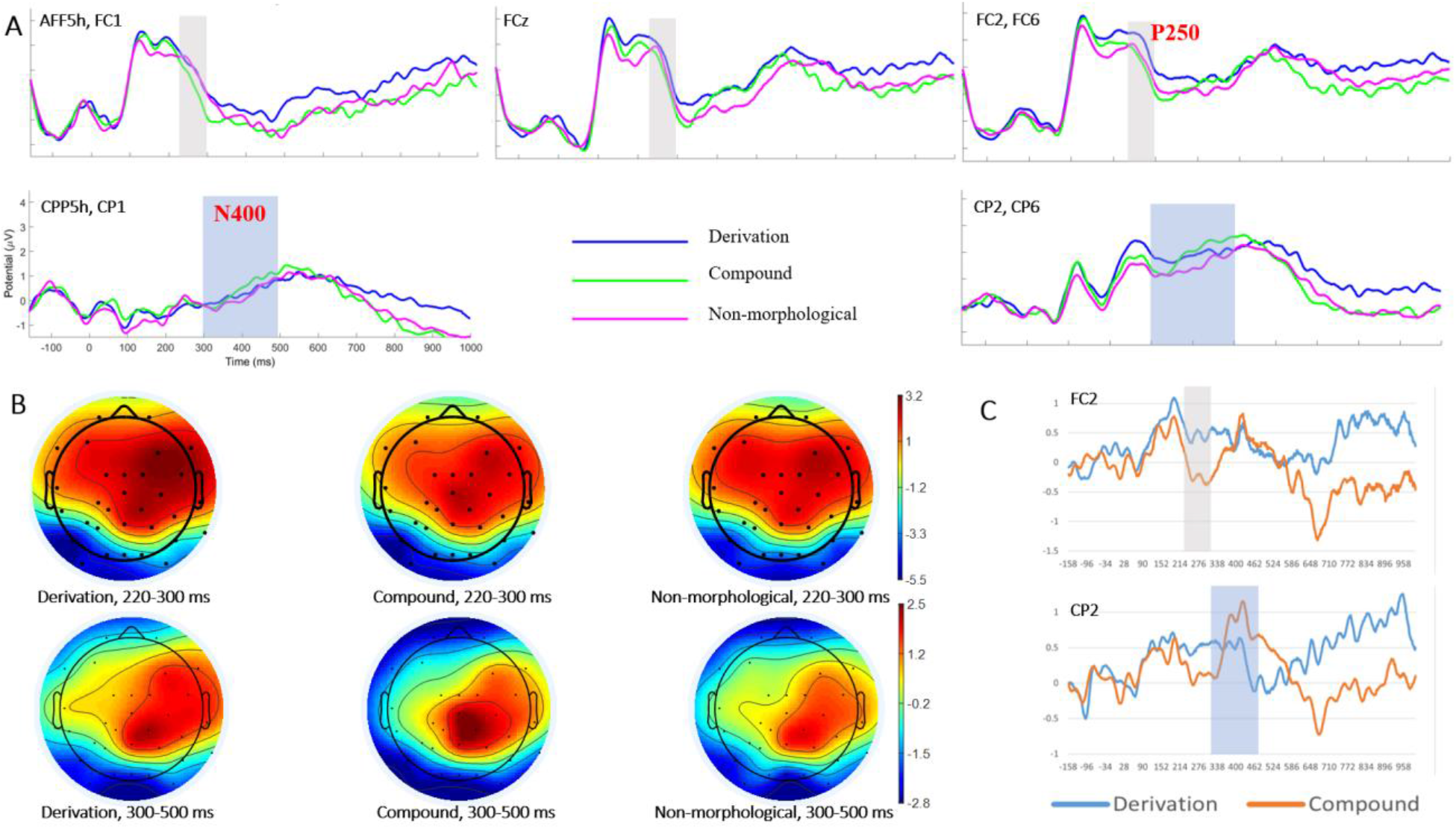
Visualization of ERP results. (A) Grand averaged ERPs over the frontal electrodes (the first row) demonstrating P250 effect (in grew shades), and centro-parietal sites (the second row) manifesting N400 activities (in blue shades). (B) Brain topographies of P250 (the first row) and N400 (the second row) across three conditions. (C) Difference waves for Derivation vs. Control (in blue), and Compound vs. Control at electrodes FC2 and CP2, respectively. The time windows of P250 and N400 were shaded in grey and blue, respectively.

**Figure 5.**
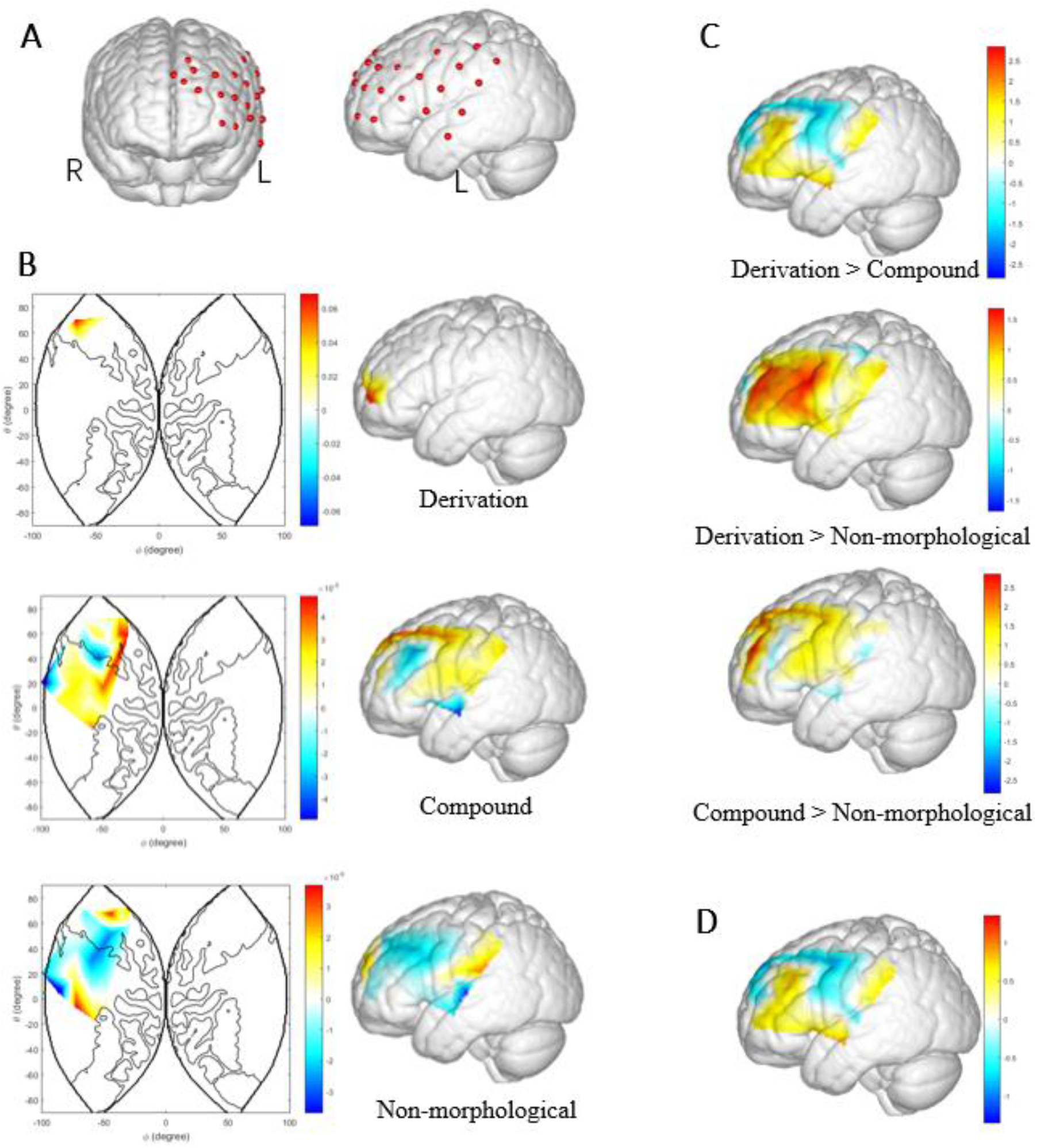
Configuration of fNIRS layout and activation maps. (A) fNIRS layout covering the frontal and temporal cortex of the left hemisphere, from the frontal and the views, respectively. (B) Brain activation patterns in reading words across three conditions based on HbO beta values. (C) T maps of three pairwise comparisons. (D) Activation difference of Derivation vs. Control (in blue), and Compound vs. Control.

The pre-processing of fNIRS data were performed with nirsLAB (Y. Xu, Graber, Schmitz, & Barbour, 2014). Data from one participate was excluded due to extensive physiology noise, leaving datasets from 29 participants for further analysis. Each fNIRS lasted 14 s, consisting of 1s before prime onset and 14 s for the hemodynamic response period. Motion artifacts were removed from the raw data by the built-in algorithm, which were subsequently filtered with a band pass of 0.01-0.2 Hz. Oxygenated hemoglobin (HbO) and deoxygenated hemoglobin concentration (HbR) changes were modeled in the Level 1 module of statistical parametric mapping with the canonical HRF function. As a result, general linear model (GLM) coefficients (beta values) were obtained across all priming conditions from each participant.

## Results

### Behavioral results

The mean accuracy rate (ACC) was 95.26%, indicating that participants well engaged the task. Repeated-measures analyses of variance (ANOVA) were conducted on reaction time (RT) and ACC, respectively. It was discovered that a significant priming effect for RT was detected, *F*(2,58)=34.085, *p*<0.001, partial *η*^2^=0.540. Multiple comparisons (Fig. 3A) revealed that RT to derivational priming (712±121ms) was significantly longer than compound constitute priming (625 ± 82ms) and non-morphological priming (636 ± 99 ms) (*p*s<0.001), whereas there was no significant difference in RT between compound and non-morphological priming (*p*>0.05).

Likewise, the priming effect was also identified for ACC, *F*(2,58)=37.873, *p*<0.01, =0.566. Derivational priming (89.87±6.19%) exhibited significantly lower ACC as compared to compound (98.8±2.31%) and non-morphological (97.1±3.03%) cases (*p*s<0.001). Meanwhile, ACC of non-morphological priming was significantly lower than that of the compound case (*p*<0.01). The behavioral results were visualized in Figure 3.

### ERP results

P250 effect was inspected during the time window of 220-300 ms after target word onset in both bilateral and midline sites of the frontal region (Chung et al., 2010). Two-way repeated-measures ANOVA was performed with priming type and hemisphere at the bilateral sites (left: AFF5h, FC1; right: FC2, FC6) as independent variables. Although no significant priming condition effect was revealed (*F*(2,58)=2.169, *p*=0.131, partial *η*^2^=0.07), the main effect of hemisphere was identified, *F*(1,29)=0.252, *p*<0.05, partial *η*^2^=0.153. In particular, significant interaction between priming type and hemisphere was discovered. In addition, the right hemisphere (2.55±0.31 μV) showed larger P250 than the left one (1.92±0.30 μV). Meanwhile, the non-morphological condition (2.10±0.33 μV) elicited significantly higher P250 than the compound case (1.54±0.37 μV) over the left hemisphere, while derivational priming (2.97±0.37 μV) evoked significantly greater P250 than the compound case (2.27±0.34 μV) along the right hemisphere. By contrast, no other comparisons reach the significance level. Likewise, the effect of priming type was examined at the midline electrode (FCz) of frontal cortex, demonstrating that no reliable difference was detected, *F*(2, 58)=1.262, *p*=0.289, partial *η*^2^=0.042.

Regarding grand-averaged ERP and topographies (Figures 4A and 4B) as well as previous studies (Chung et al., 2010), N400 effect was examined in the time windows of 300-500 ms with priming categories and hemispheres (left: CPP5h, CP1; right: CP2, CP6) as factors. Significantly lower N400 were detected in the left hemisphere (0.295±0.268 μV) than that in the right one (1.756±0.278 μV), *F*(1,29)=40.433, *p*<0.001, partial *η*^2^=0.582. No other significant main effect or interaction was detected.

To distinguish well between the three different word structures, the priming effect was accessed by comparing the difference ERP waves between the two priming conditions, namely, the P250 and N400 amplitudes of compound minus that of control and derivation minus control, respectively (Figure 4C). Derivational structures (derivation minus control, 0.279±0.350 μV) elicited greater P250 than compounding structures (−0.354±0.250 μV) (*p*s<0.05) in both hemispheres. Interestingly, P250 in the right hemisphere (0.206±0.294 μV) was statistically higher than that from the left hemisphere (−0.280±0.261 μV) (*p*<0.05). The P250 patterns in the midline were consistent across the bilateral sites. With regards to N400 component, the patterns remained unchanged.

### fNIRS results

Although both HbO and HbR beta values were generated from the GLM estimations, the present study only analyzed the HbO signals to examine the priming condition effect (He, Wang, Li, & Yuan, 2017; Hu et al., 2018). Firstly, HbO beta values were compared across the three conditions by repeated-measures ANOVA channel by channel, among which channel 6 in the frontopolar area showed a significant condition effect, *F*(2, 56)=3.162, *p*<0.05, partial *η*^2^=0.101. Pairwise comparisons were then conducted between Derivation vs. Compound, Derivation vs. Non-morphological, and Compound vs. Non-morphological conditions. Table 2 summarized significant (*p*<0.05) and marginal significant (0.05<*p*<0.1) results when *p* values were not corrected. Specifically, the left frontal cortex including the pre-motor and supplementary motor cortex (channels 5, 6, 8. 11, and 18) exhibited enhanced activation from non-morphological condition to compound words. Meanwhile, compound produced larger frontal cortex activation than derivations, as manifested in channel 10. In particular, the associated brain activation patterns were visualized in Figures 5B and 5C.

**Table 2.**
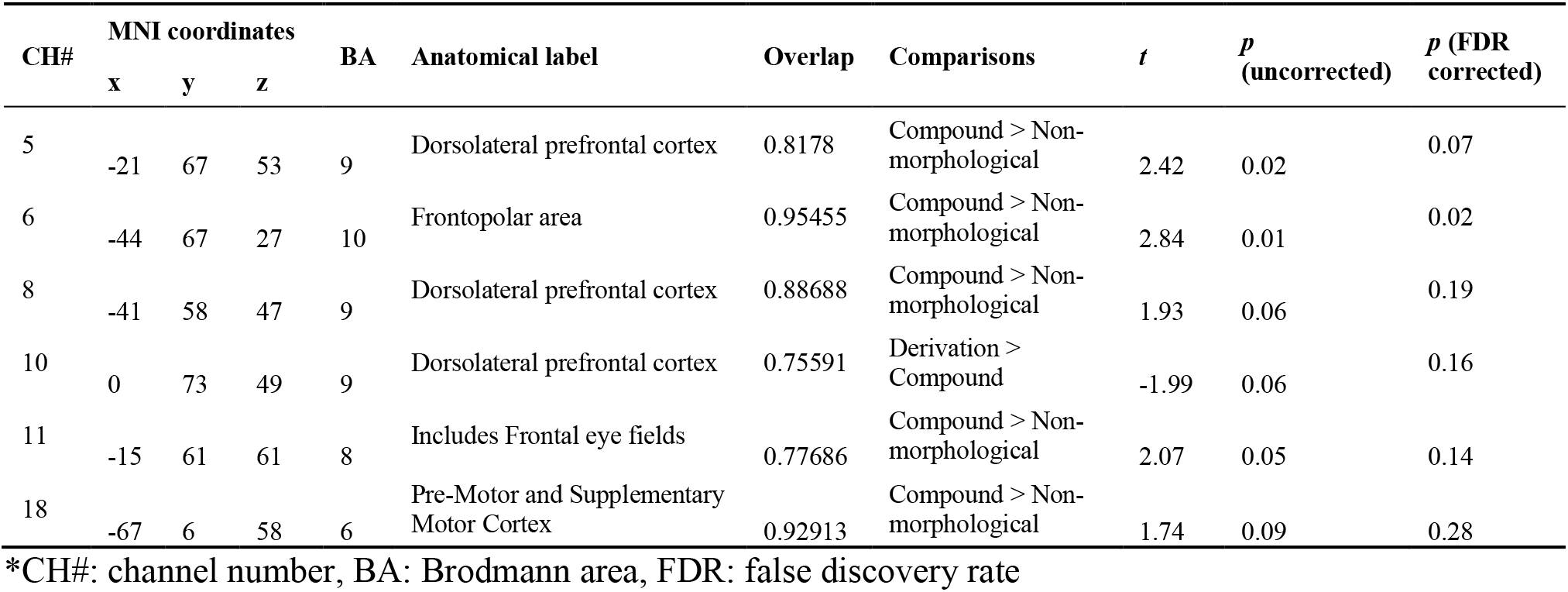
Comparison results of significant and marginal significant channels and their corresponding spatial information.

Besides, pure word structure effect (derivation/compound minus non-morphological) was also examined and the *t* tests results were plotted in Figure 5D. It was discovered that only channel 10 illustrated a marginal significance (*p*=0.057), in which compounds exhibited enhanced brain activation than derivations.

### Correlational results

The relationships between P250 and HbO beta weights were examined by calculating the Pearson correlation coefficients. Specifically, mean ERP amplitudes of FP1, F7, AFF5h, FC1, FCz and fNIRS hemodynamic responses on channels 5, 6, 8, 10 and 11 were used to depict brain activation in the left frontal cortex, respectively, while the temporal responses were accessed from EEG channels (FTT7h, TTP7h, CPP5h, CPP3h, CP1) and fNIRS channels (CH15, CH16, CH17, CH21, CH22). According to the matrixes of correlation coefficients, P250 well predicted the temporal activation in derivation reading (Figures 6A and 6B), which implicates a broad semantic network. More importantly, both spatial (frontal) and temporal ROIs (Figures 6C and 6D) showed close associations between P250 and hemodynamic responses in derivation vs. compound contrast.

**Figure 6.**
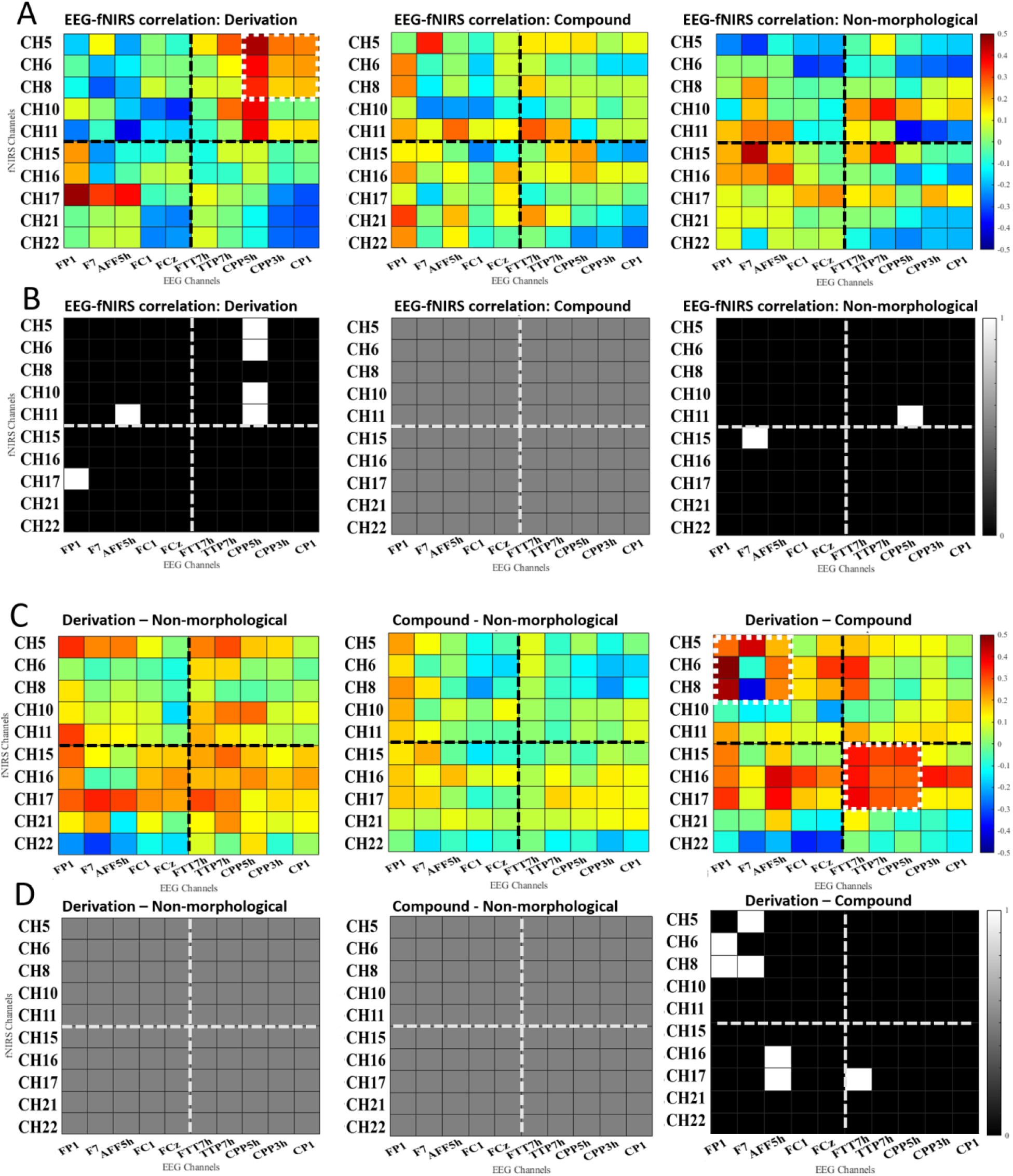
Correlation coefficient matrixes between electrophysiological responses (ERP data) and brain activation (fNIRS data). Dashed line represents the division of frontal (ERP: FP1, F7, AFF5h, FC1, FCz; fNIRS: CH5, CH6, CH8, CH10, CH11) and temporal regions (ERP: FTT7h, TTP7h, CPP5h, CPP3h, CP1; fNIRS: CH15, CH16, CH17, CH21, CH22). (A) Correlations between ERP and fNIRS data among representative probes across three conditions. Brighter color stands for stronger correlation. The white square with dashed outline represents region of interest. (B) Binary maps of the correlation results across three conditions. The white squares denote a significant correlation between bi-modal data (*p*<0.05). (C) Correlations between ERP and fNIRS data among representative probes across three comparisons. (D) Binary maps of the correlation results across three comparisons (*p*<0.05).

## Discussion

The current study examined both morphological priming effect (compound/derivation constitute priming vs. non-morphological priming) and word structure effect (derivation vs. compound) by using a masked priming technique in a lexical decision task. The analysis of behavioral performance revealed a morphological structure facilitation for coordinate compounds, which exhibited statistically higher accuracy and relatively shorter reaction time than non-morphological pairs. This pattern extended the findings from Chung et al. (2010) by suggesting that coordinate structure priming effect could not only be elicited from word pairs sharing the same structure, but also in coordinate words which are primed by their first roots, conditional on a short SOA (57 ms). Nevertheless, this morphological facilitation was not in line with P. D. Liu and McBride-Chang (2010a) and Cui et al. (2018). The discrepancy could be attributed to the relatively longer SOA and different task demands, where the former used 200 ms for SOA and the latter asked participants to respond to both primes and targets in a self-paced manner. The current findings therefore implicate that Chinese coordinate structure effect might be sensitive to SOAs and index an automatic and short-lived activation of morphological information at an early stage of lexical access (P. D. Liu & McBride-Chang, 2010a). Yet, the present study failed to detect any morphological priming effect for Chinese derivations. As there is no previous study examining the psychological reality of Chinese derivations, and morphological structure effect is somehow weak in behavioral patterns due to technical limitation (Chung et al., 2010; Cui et al., 2018), we will rely more on ERP and imaging data to advance understandings into this word structure.

With respect to word structure effect, both reaction time and accuracy data revealed that derivations might be harder to recognize than coordinate structures in constitute priming conditions. This finding is somewhat consistent with the coordinate vs. subordinate structure contrast identified in semantic priming paradigm (P. D. Liu & McBride-Chang, 2010a). Both subordinate structure (e.g., *黑板*, black-board, “blackboard”) and derivation (e.g., *作家*, writingexpert, “writer”) employed a modifier-head relation, whereas the suffix in derivation (i.e., *家* in *作家*) has been delexicalized in grammaticalization process. As a result, the suffix is loosely attached to the base form and productive in deriving new word forms (e.g., *画家*, “painter”; *艺术家*, “artist”; *音乐家*, “musician”). In contrast, the inter-connection between the constitutes within a coordinate structure is relatively stronger than the base-suffix association in a derivation, even not as strong as those of subordinate compounds (D. Liu, 2016). Therefore, the word structure effect identified in the current behavioral results shed light on the spreading activation account, where the strengths between constitutes within a morphological complex word would determine the cognitive efforts needed in word recognition.

The ERP data revealed a prominent word structure effect on the frontal P250. Difference wave analyses showed that derivations elicited significantly greater bilateral negativities in the time window of 220-300 ms, compared to coordinate compounds, while the right hemisphere tends to be more significant. The frontal P250 effect was previously found in the comparison between word pairs of different semantic relatedness (Hill et al., 2002) and structure consistency (Chung et al., 2010). Importantly, this early component was sensitive to SOA, as it was only present in a short SOA of 150 ms/57 ms, yet not in 700-ms condition, which indexes the automatic access in lexical processing. Existing MEG studies (Cavalli et al., 2016) also associated this early time window with a decomposition and parsing operation in morphological processing. As such, relative to coordinate compounds, greater P250 amplitudes found in derivations of the current study would indicate more cognitive resource consumption in morphological parsing, which confirmed the behavioral patterns. By virtue of the spreading activation account, all concepts (i.e., word roots in the current case) are stored as nodes and connected with each other by different weights. With the input of prime, not only the corresponding morphemes but also the candidate morpheme, with which it could make up new words, would be activated, followed by the recombination of two morphemes. Yet, the inter-constitute connection across word structures is different. For instance, compared to subordinate structures, both constitutes in coordinate structure contribute equally to the whole word, manifesting a rather loose connection (D. Liu, 2016). The current P250 effects further suggest that Chinese derivations and compounds could be discriminated in the frontal cortex in light of their differing constitute relationships, as facilitated by spreading activation.

In addition to P250 effects, Chung et al. (2010) also reported a classic N400 semantic priming effect mediated by semantic relatedness between primes and targets, which was absent in the current study. According to the grand-average brainwaves and topographies, there was a N400-like component, which was widely distributed in the centro-parietal cortex. Yet, the comparison between three conditions did not reach any statistical difference. It was because the current study did not manipulate the semantics, as the semantic associations between primes and targets were generally comparable by measuring and matching semantic transparency. The frontal P250 effect could therefore be attributed to a relatively purer word structure modulation.

Brain activation obtained from fNIRS data provided robust evidence on morphological priming effect in compound word recognition, which was not statistically reliable in reaction time. Coordinate compounds elicited enhanced activation than the controls in the left frontal network, including dorsolateral prefrontal cortex (DLPFC) and frontopolar area. This pattern identified in the left frontal cortex from the current study is roughly consistent with previous fMRI results associated with morphological processing for children and/or with auditory stimuli. For instance, word pairs sharing morphemic information (e.g., *高温-高空*, high temperature-high sky) showed the highest activation in the left inferior frontal gyrus (IFG) across all lexical conditions in an explicit auditory morphological judgment task (Zou et al., 2015). This implicates an important role of the left IFG in morphological processing across alphabetic languages (Bozic, Marslen-Wilson, Stamatakis, Davis, & Tyler, 2007; Lehtonen, Vorobyev, Hugdahl, Tuokkola, & Laine, 2006) and Chinese. In a recent bilingual children study (Ip et al., 2017), Chinese-English bilingual children elicited greater activation in the IFG in auditory English morphological task (e.g., re-jump), compared to Chinese compound morphology task (e.g., *病花*, sick-flower) This pattern was similar to monolingual English children, further highlighting the role of left frontal cortex in morphological processing. Compared to fMRI, fNIRS is relatively limited in spatial resolution, which might cause the current study’s failure to localize subtle changes in exact IFG. Yet, the left frontal cortex (mostly DLPFC) still showed manifested activations in morphological priming conditions. Importantly, the current findings could justify previous fMRI results with data from print reading among adult population. Taken together, we may infer that the left frontal cortex implicates a core area for morphological processing across language modalities and literacy stages. Whilst this study did not include as many word structures, it did partially substantiate a word structure effect between Chinese derivations and coordinate compounds, which made an original contribution to the field. Even though the contrast between the two structures only reached a statistically marginal significance, correlational analysis on derivation vs. compound difference revealed a good coherence between frontal P250 values and hemodynamic responses in the left frontal area. Derivations generated stronger activation than compounds, given the rather loose connections between roots and suffixes, which might demand more efforts in constitute meaning access and re-composition. The fNIRS data is overall consistent with ERP results and further validate the word structure effect in the frontal cortex.

Nevertheless, unlike previous MEG and fMRI studies (Hsu et al., 2019; Ip et al., 2019), the current study did not detect exact temporal engagement neither in morphological processing nor in word structure differentiation. Yet, ERP-fNIRS correlations showed that P250 in temporal cortex could well predict brain activations in the corresponding regions. Meanwhile, the electrophysiological responses in the left temporal region were closely associated with frontal activations. Whilst we highlighted the role of the left frontal cortex in word structure discrimination, the temporal areas should not be neglected. Future studies could address this region by using fMRI technique and functional connectivity analysis.

Finally, the current findings did not apply to the dual-route theory in morphological rich language (Baayen et al., 1997; Bozic & Marslen-Wilson, 2010). According to this theory, the recognition of regular inflections and rule-based word units requires online computation mechanism and employ the left fronto-temporal cortex. In contrast, storage-based and highly lexicalized items engage broader bilateral brain regions, which is less time-costly. However, Chinese is a language lacking morphological inflection. Instead, compounding is the dominant structure of Chinese morphology and implicates a storage-based representation. Even though the proportion of derivations is much smaller than compounds, they are highly productive given the adhesiveness of affixes. By comparing the two word structures with differing lexicalization extents, the current study found that Chinese derivations might employ broader brain networks in bilateral fronto-temporal cortex, while compounds were mostly manifested in the left frontal regions and indexes a storage-based mechanism. This brain pattern might be attributable to the connection strength between constitutes across different word structures in light of spreading activation theory.

Together, this study examined both morphological priming effect and word structure effect by using EEG-fNINS simultaneous data. Specifically, we found prominent word structure effect in the frontal cortex. Chinese derivations elicited greater activation in this area and engaged more distributed network than lexicalized compounds, as manifested by both electrophysiological and hemodynamic responses. As a morphologically impoverished language, Chinese word structure effect showed a distinct pattern from the dual-route mechanism in alphabetic languages. Future research needs to be done to examine this hypothesis by including more morphological structures and using more nuanced imaging technique.

## Data Availability Statement

Data is available on request.

## Acknowledgment

This work was supported by the University of Macau (MYRG 2020-00067-FHS, MYRG2019-00082-FHS and MYRG2018-00081-FHS), the Macao Science and Technology Development Fund (FDCT 0020/2019/AMJ and FDCT 0011/2018/A1), and Higher Education Fund of Macao SAR Government (CP-UMAC-2020-01).

